# Cross-species prediction of transcription factor binding by adversarial training of a novel nucleotide-level deep neural network

**DOI:** 10.1101/2024.02.06.579242

**Authors:** Qinhu Zhang

## Abstract

Cross-species prediction of TF binding remains a major challenge due to the rapid evolutionary turnover of individual TF binding sites, resulting in cross-species predictive performance being consistently worse than within-species performance. In this study, we first propose a novel Nucleotide-Level Deep Neural Network (NLDNN) to predict TF binding within or across species. NLDNN regards the task of TF binding prediction as a nucleotide-level regression task. Beyond predictive performance, we also assess model performance by locating potential TF binding regions, discriminating TF-specific single-nucleotide polymorphisms (SNPs), and identifying causal disease-associated SNPs. Then, we design a dual-path framework for adversarial training of NLDNN to further improve the cross-species prediction performance by pulling the domain space of human and mouse species closer.

## Background

Transcription factors (TFs) play an essential role in transcriptional regulatory networks by binding to non-coding functional regions at specific locations to promote or repress the transcription of target genes[1]. GWAS (Genome-wide association study) has revealed that the vast majority of mutations in the human genome exist in non-coding functional regions, and some of them may directly disrupt DNA binding affinities of TFs, sequentially altering the regulation of gene expression and causing some specific diseases[2]. Identification of TF binding, therefore, is always a fundamental step for molecular and cellular biology.

In recent years, deep learning (DL) has achieved considerable success in predicting epigenomic profiles from DNA sequences, including transcription factor binding[3-5], chromatin accessibility[6, 7], and histone marks[8, 9]. By learning a sequence-function relationship, trained DL-based models have been utilized on various downstream tasks, such as predicting the functional effects of single-nucleotide variants associated with human diseases[10]. These DL-based models are roughly divided into two categories: sequence-level models and nucleotide-level models. Sequence-level models take sequences as input and output scalars representing label probabilities. For example, DeepBind[3] applied a convolutional neural network (CNN) to predict the sequence specificity of TF binding. DeepSea[6] similarly used a deeper CNN to predict TF binding motifs and the chromatin effects of sequence alterations from large-scale chromatin-profiling data. DanQ[11] applied a hybrid network combining CNN with recurrent neural network (RNN) to predict TF binding motifs and prioritize functional SNPs. In contrast, nucleotide-level models take sequences as input and output vectors recording the label probabilities of nucleotides. For instance, FCNA[12] proposed a fully convolutional neural network (FCN) to predict TF binding and discovery motifs on ChIP-seq data. Leopard[13] applied an encoder-decoder architecture to predict cross-cell-type TF binding sites at the nucleotide level by integrating DNA sequences and chromatin accessibility data. BPNet[14] developed a dilated CNN to predict base-resolution ChIP-nexus binding profiles of pluripotency TFs from DNA sequences. FCNsignal[15] designed a symmetrical encoder-decoder architecture to predict base-resolution ChIP-seq binding signals of TFs from DNA sequences. However, an open question of how to compare the two types of DL-based methods also arises in a unified framework. To answer this question, Kelley *et al*.[16] proposed to ‘binarize’ their quantitative predictions, enabling a comparison of the overlap with binary labels. Avsec *et al*.[14] compared the performance of a binary model with an augmented version that appends an output head that simultaneously predicts quantitative profiles. Toneyan *et al*.[17] designed a unified evaluation framework to compare various binary and quantitative models by calculating the average coverage predictions at positive regions and negative regions based on corresponding binary-labeled data. Nonetheless, these methods require extra effort to deal with outputs or retrain models, which may be sensitive to hyper-parameter choice and parameter initialization. Therefore, it remains challenging to make a direct comparison of the two types of DL-based models using standard metrics.

Recent studies have revealed that gene expression patterns and TF binding preferences are broadly conserved across closely related species[18, 19]. Since the amino acid sequences of TF proteins, their DNA binding domains, and intrinsic DNA sequence preferences are typically highly conserved, the sequence preferences of TF binding in one species are predictive of those in closely related species. Accordingly, several computational approaches have been proposed to demonstrate the feasibility of cross-species prediction of regulatory profiles[20-22]. However, cross-species TF binding prediction is complicated by the rapid evolutionary turnover of individual TF binding sites across the genomes of different species, even within cell types that have similar functions. Cochran et al.[23] conducted a comprehensive analysis to show that cross-species predictive performance is consistently worse than within-species predictive performance and proved that this difference is partly caused by species-specific repeats; in addition, they developed a domain-adaptive method for improving overall cross-species model performance. Although this method works well for cross-species prediction of four TFs in the liver, it fails to generalize well to an expanded set of TFs from different cell lines. Moreover, most approaches merely focus on the feasibility of cross-species TF prediction; however, direct training of cross-species models for TF binding prediction has been less explored.

In this study, we mainly focus on two unresolved issues mentioned above: (i) there is still a lack of an effective method for benchmarking sequence-level models versus nucleotide-level models directly; (ii) there is an urgent need for an effective method to directly train cross-species models for TF binding prediction. On the one hand, inspired by the concept of pixel-level image segmentation[24], we develop a novel Nucleotide-Level Deep Neural Network (NLDNN) to predict TF binding within or across species. NLDNN consists of an encoder, a decoder, and a connection, forming a symmetrical ‘U’ shape architecture, which takes DNA sequences as input and directly predicts experimental coverage values. Furthermore, we propose to use the maximums of coverage values as a bridge between sequence-level and nucleotide-level models, so as to make a direct comparison of the two types of models. On the other hand, we design a dual-path framework for adversarial training of NLDNN to reduce the cross-species prediction performance gap by pulling the domain space of different species closer. Resembling GAN, the framework consists of two generators, one for source species and another for target species, a predictor, and a discriminator. The experimental results show that NLDNN performs better than competing methods (including sequence-level and nucleotide-level models) in multiple prediction tasks and that adversarial training can improve the overall performance of cross-species TF binding prediction.

## Results

### Overview

To study TF binding at the nucleotide-resolution level, we propose a Nucleotide-Level Deep Neural Network (NLDNN), as shown in Fig.1a-b, which takes DNA sequences as input and directly predicts coverage values. Unlike binary models, which regard TF binding prediction as a sequence-level classification task, NLDNN regards it as a nucleotide-level regression task. Considering that high coverage values are most likely enriched in TF binding regions, we, therefore, propose to use the maximums of coverage values to represent the strength of TF binding, by which binding regions are easily distinguished from non-binding regions since the strength of binding regions is generally higher than that of negative regions. To verify the rationality of this assumption, we computed the Pearson correlation between the maximums of coverage values and the strength of TF binding (see Methods for details), high average values (human: 0.9; mouse: 0.818) showing that the maximums of coverage values are closely related to the binding strength (Supplementary Fig.1). Through this way, we can make a direct comparison of sequence-level and nucleotide-level models by evaluating the classification and fitting ability of them using some standard metrics (Fig.2a), e.g. PR-AUC, Pearson correlation. Beyond predictive performance, we also assess the ability of NLDNN to locate potential TF binding regions and predict variant effect and delve into the nucleotide’s contribution to TF binding or variant prediction via model interpretability. To further improve the cross-species predictive performance of NLDNN, we design a dual-path framework (Fig.1e) in which the top path is used to generate source feature mappings, the bottom path is used to generate target feature mappings and the discriminator is used to classify species labels. Next, we perform adversarial training to learn a target generator such that it can learn a shared feature space between species.

**Fig. 1.**
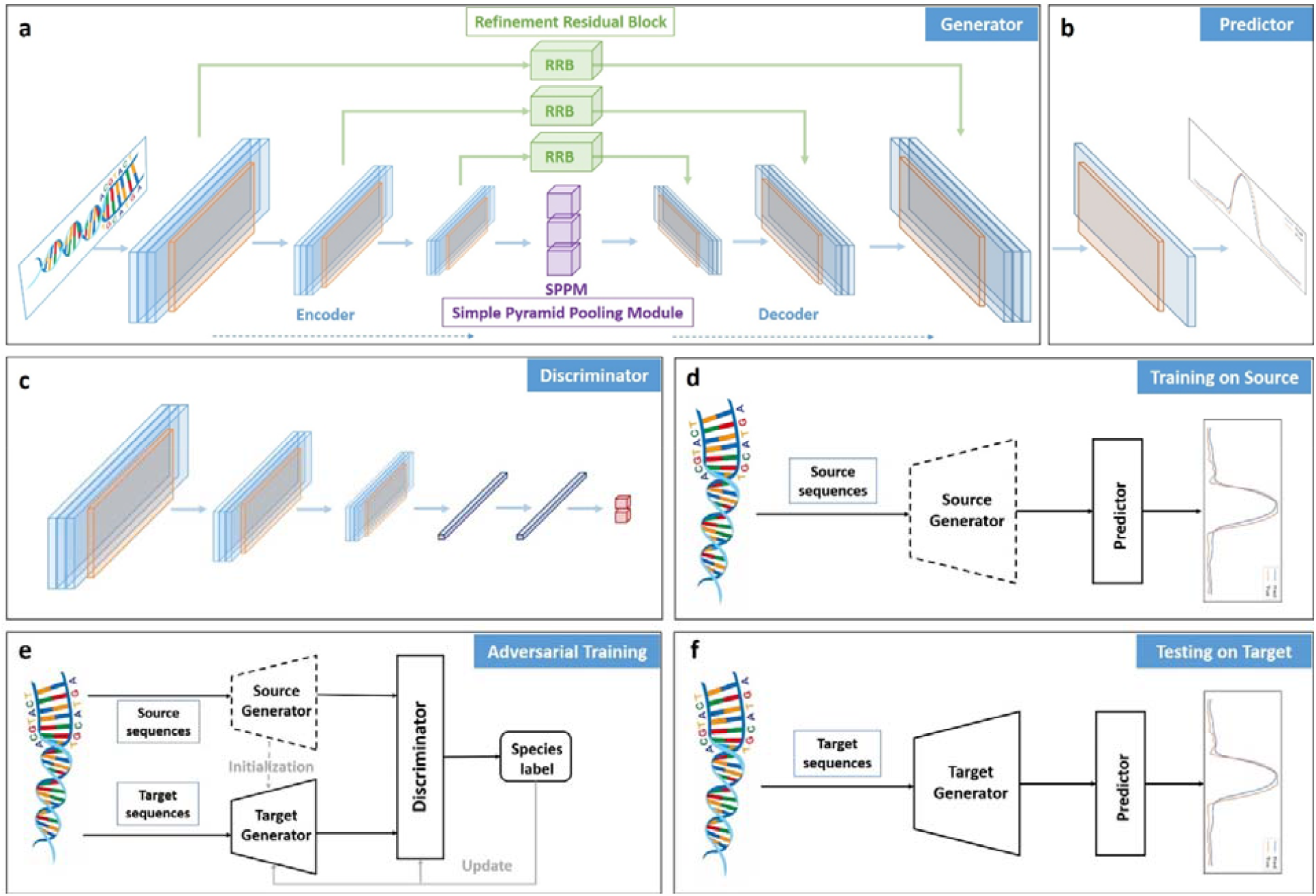
The flowchart of our proposed method (NLDNN-AT). (a-b) The architecture of NLDNN can be decomposed into a generator (a) and a predictor (b) during adversarial training. (c) The architecture of a discriminator. (d) The framework of training NLDNN on source species. (e) The framework of adversarial training is a dual-path architecture consisting of a source generator, a target generator, and a discriminator. (f) The framework of testing NLDNN on target species. The whole process is divided into three stages: (i) in the training stage, we trained a source generator and a predictor using data from a source species; (ii) in the adversarial training stage, we first used the parameters of the source generator to initialize a target generator and then applied adversarial training to fine-tune the target generator; (iii) in the testing stage, we used the target generator and predictor to test data from a target species.

### Classification vs. Regression models

This study raises a question: classification and regression models, which one is better for predicting TF binding? To explore this question, two types of sequence-level models from competing methods, including classification and regression models, were used. Sequence-level classification models take DNA sequences as input and predict their corresponding label probabilities ranging from 0 to 1. On the contrary, we remodified three classification models (DeepSea, DanQ, DanQV) as corresponding regression models (DeepSea+, DanQ+, DanQV+) by removing the last sigmoid layer from them and replacing true labels with the maximums of experimental coverage values. Regression models, therefore, take DNA sequences as input and predict the maximums of coverage values. The classification performance of these models can be evaluated directly using the PR-AUC metric. After removing the last sigmoid layer from classification models, their fitting performance can be compared fairly using the Pearson correlation metric(Fig.2a).

**Fig. 2.**
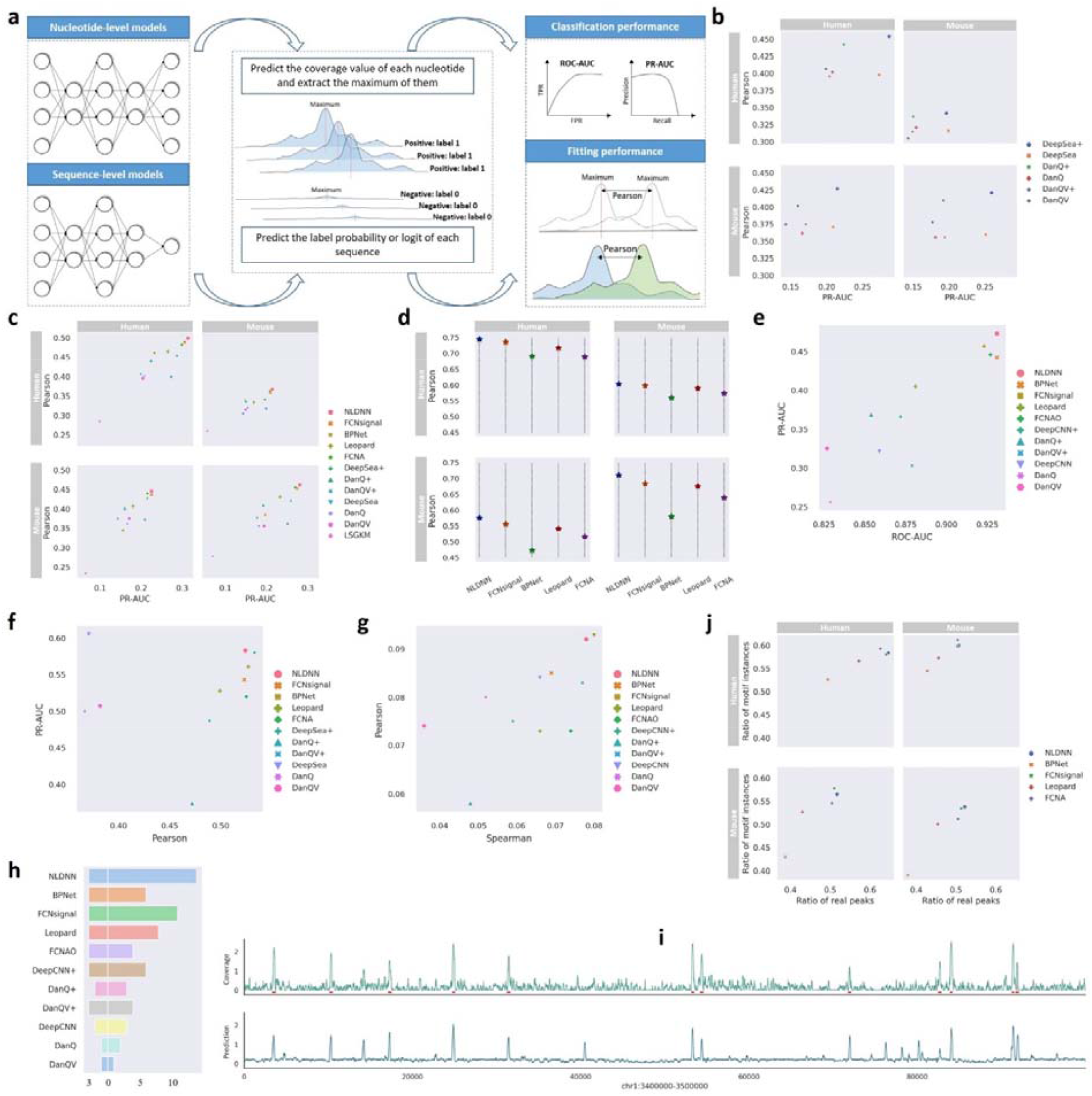
Evaluation of sequence-level and nucleotide-level models. (a) Schematic overview of directly comparing sequence-level and nucleotide-level models. (b) Within- and cross-species performance comparison between sequence-level classification and regression models for predicting TF binding. (c) Within- and cross-species performance comparison between all models for predicting TF binding. (d) Pearson correlation of all nucleotides across nucleotide-level models. (e) Performance comparison between all models for classifying TF-specific SNPs filtered by peaks. (f) Performance comparison between all models for predicting chromatin accessibility signals. (g) Performance comparison between all models for fitting MPRA data. (h) Performance comparison between all methods for prioritizing causal SNPs. (i) Visualization of the true and predicted coverage values of a human genomic region (chr1:3400000-3500000, 100kb), where red underlines denote true peaks on the genomic region. (j) Within- and cross-species performance comparison between nucleotide-level models for locating TF binding regions.

As shown in Fig.2b, regarding the fitting performance (Pearson correlation), DeepSea+ and DanQ+ perform much better than DeepSea and DanQ for within- and cross-species prediction, whereas DanQV+ performs better than DanQV for within-species prediction but worse for cross-species prediction. Regarding the classification performance (PR-AUC), DeepSea+ and DanQ+ perform better than DeepSea and DanQ for within-species prediction but worse for cross-species prediction, whereas DanQV+ performs worse than DanQV for within- and cross-species prediction. Overall, the results show that sequence-level regression models are better than sequence-level classification models with performance gains of 0.2% (PR-AUC) and 3% (Pearson correlation), demonstrating that regression models have advantages over classification models in terms of fitting and classification performance.

### The overall performance of NLDNN

The previous section mainly investigated the performance of classification and regression models from a sequence-level perspective. In this section, we intend to make a comprehensive comparison of sequence-level and nucleotide-level models. To evaluate the within- and cross-species performance of NLDNN, several competing methods were used, including seven sequence-level models and four nucleotide-level models. Beyond directly assessing predictive performance, we assess each model with additional criteria: (i) variant effect prediction and (ii) localization ability (see Methods for details).

#### Predictive performance

To comprehensively compare the predictive performance of NLDNN with that of competing methods, the PR-AUC and Pearson correlation were adopted to separately assess their classification and fitting performance. Through the maximums of coverage values, the predictive performance of all methods can be directly compared. From the experimental results (Fig.2c and Supplementary Fig.2), we have the following observations: (i) nucleotide-level models are generally better than sequence-level models for within- and cross-species TF binding prediction; (ii) among nucleotide-level models, NLDNN performs best in terms of the classification (PR-AUC) and fitting (Pearson correlation) performance, demonstrating the effectiveness of NLDNN for TF binding prediction; (iii) compared to within-species prediction, the performance of cross-species prediction is decreased by a large margin, indicating that the task of cross-species prediction is more complex. In addition, we calculated the Pearson correlation of all nucleotides for nucleotide-level models (Fig.2d), further confirming the superiority of NLDNN over other nucleotide-level models. Overall, the above results prove that the predictive performance of NLDNN is better than that of the competing methods in terms of classification and fitting ability.

#### Variant effect prediction

To assess the performance of NLDNN and competing methods for variant effect prediction, we constructed three tasks: classification, regression, and prioritization. For classifying TF-specific SNPs, the ROC-AUC and PR-AUC were adopted to assess the classification performance of all methods. As shown in Fig.2e, we find that (i) nucleotide-level models are better than sequence-level models; (ii) sequence-level regression models perform better than sequence-level classification models; (iii) NLDNN performs best among all methods in this task. For fitting MPRA data, we first need to prepare trained models using ATAC-seq or DNase-seq data, which are known as chromatin accessibility prediction tasks, and the comparative results of all methods on this task are visualized in Fig.2f. Interestingly, DeepSea performs best among all methods in terms of classification performance, and DeepSea+ performs best among all methods in terms of fitting performance. These observations are inconsistent with those found in the task of TF binding prediction, suggesting that nucleotide-level models are better suited for TF binding prediction. The underlying reason may be that nucleotide-level models are good at mining TF-specific motif features while sequence-level models are good at integrating the contextual information of a whole sequence. Nevertheless, NLDNN has comparable performance to DeepSea and DeepSea+. Then, Pearson and Spearman correlations were adopted to assess the fitting performance of all methods on MPRA data. As shown in Fig.2g, we observe that (i) nucleotide-level models are generally better than sequence-level models; (ii) FCNsignal performs best among all methods, followed by NLDNN. For prioritizing causal SNPs, we counted the number of real SNPs identified from three disease-associated SNP groups (left panel in Fig.2h) and the ratio of the effect values of real SNPs to the highest values of SNPs with strong LD (right panel in Fig.2h). Evidently, nucleotide-level models successfully identify the causal SNPs in myeloma (rs4487645), CLL (rs539846), and pan-autoimmune (rs6927172), while most of sequence-level models fail to identify all of them, and NLDNN has the highest difference ratio. Overall, the performance of NLDNN is better than that of the competing methods on the tasks of variant effect prediction.

#### Localization ability

Since nucleotide-level models are able to predict the coverage values of nucleotides, they all have the ability to locate potential TF binding regions on chromosomes. Although all models were trained using DNA sequences of length 600bp, the trained models can accept inputs of arbitrary length and predict each nucleotide’s value (Fig.2i). To assess the performance of nucleotide-level models for locating TF binding regions, we counted the number of all located regions intersecting with true peaks and the number of found motif instances (Methods). As shown in Fig.2j, the localization ability of NLDNN is better than that of other nucleotide-level models according to the two evaluation ways, particularly the ability to identify real peaks from genomes.

Based on the three criteria, the overall performance of NLDNN is better than that of the competing methods, demonstrating the effectiveness of NLDNN for TF binding prediction.

### Model interpretation

#### Motif discovery

A primary downstream application of DL-based models is the discovery of functional motifs and their complex interactions. Trained models can provide a direct way to learn translational patterns, such as motifs through the first convolutional layer, whose kernels are regarded as motif scanners. In this way, NLDNN, DeepSea+, and DeepSea were used for motif discovery. Besides, TF-Modisco was also applied to do motif discovery, which can discover TF motifs from importance scores profiled by DeepLift. From the results (Fig.3a), we find that improved predictions may not necessarily lead to learning better motifs; as you see, the matching significance of NLDNN is slightly worse than that of DeepSea+ and DeepSea; however, the motifs learned by the model itself are better matched with known motifs than those learned by TF-Modisco. The decreased performance of TF-Modisco may be caused by insufficient data as we used only positive sequences from the test set. Irrespective of how the task was framed, the vast majority of known motifs and co-binding motifs have been successfully identified (Fig.3b). For instance, the co-binding motifs of JUND were mainly Fos-related motifs (e.g., FOS, FOSL1 and FOSL2) and ATF-related motifs (e.g., ATF2, ATF3), supported by the evidence that the activator protein 1 (AP-1) is assembled from JUN-JUN, JUN-FOS, or JUN-ATF family protein homo- or heterodimers to transactivates or represses its target genes[25]. The co-binding motifs of GATA1 contain JUN and FOS with significant *q*-value, supported by the evidence that GATA family frequently occupies the same chromatin sites as c-JUN and c-FOS, heterodimeric components of AP-1[26].

**Fig. 3.**
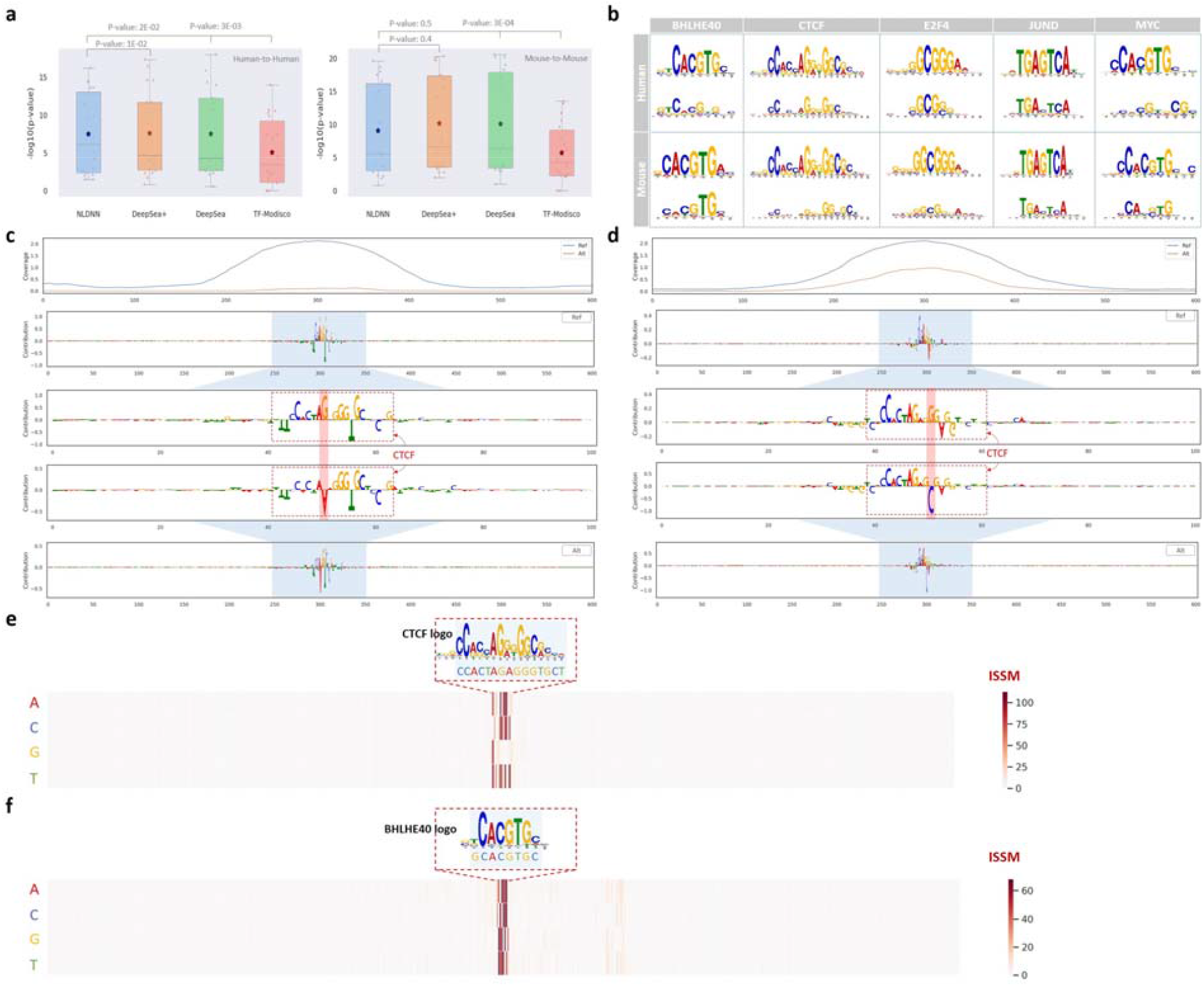
Motif discovery and interpretation of SNP. (a) The matching degree (-log(*p*-value)) of TF motifs learned from different models within species. (b) Visualization of five learned TF motifs within species. (c) Visualization of a single nucleotide variant (G→A) within CTCF binding sites, significantly affecting TF binding. (d) Visualization of a single nucleotide variant (G→C) within CTCF binding sites, slightly affecting TF binding. (e) Visualization of important sites of a CTCF binding sequence identified through ISSM, corresponding precisely to the CTCT motif. (f) Visualization of important sites of a BHLHE40 binding sequence identified through ISSM, corresponding exactly to the BHLHE40 motif.

#### Interpretation of SNPs

As the disruption of TF binding affinities may alter the expression of genes associated with diseases, figuring out how single nucleotide variants affect the binding of TFs is a crucial step for studying the pathogenic mechanisms of diseases. To achieve this goal, taking TF-specific SNPs as examples, we first applied DeepLift to compute the contributions of nucleotides in reference and altered sequences, respectively. Then, we investigated how single-nucleotide variants influence TF binding activity by observing the changes from reference sequences to altered sequences. By visualizing the changes between them, we find that some types of single-nucleotide variants may dramatically influence TF binding although the motif logos of TFs remain basically unchanged, e.g., single-nucleotide variants (G→A, G→T) within CTCF binding sites almost reduce the coverage values to zero (Fig.3c and Supplementary Fig.3a), implying that the nucleotide G at the mutated position is very important for CTCF binding. However, a variant (G→C) within CTCF binding sites just reduces the coverage values by a small margin, perhaps because G and C are a complementary base pair (Fig.3d). We also find that some variants do not play a major role in affecting TF binding instead dominated by its co-binding factors, e.g., a variant (A→G) within E2F4 binding sites has little effect on TF binding even though the contributions of this binding region are vanished (Supplementary Fig.3b), implying that TF binding to this region is dominated by its upstream co-binding factor (E2F family). More examples are displayed in Supplementary Fig.4. Through interpretation of SNPs, we find that single-nucleotide variants within binding sites indeed affect TF binding in various ways.

To further investigate the importance of each nucleotide within TF binding sites, we applied ISSM to calculate the effect values of variants at each position in binding sequences. By visualizing examples of CTCF and BHLHE40 binding regions (Fig.3e-f), we observe that these important variants with high ISSM values correspond precisely to their binding sites (motif logo).

For instance, the identified sites ‘CCACTAGAGGGTGCT’ are a part of the CTCF motif logo and necessary for TF binding, as changing any of these nucleotides will result in significant changes in predictive performance (high ISSM values). Significantly, the changes of C or G at positions 5, 10, and 13 in the CTCF motif logo make TF binding prediction much worse, indicating that these conservative sites are more crucial for CTCF binding. More examples with similar results are displayed in Supplementary Fig.5.

#### Interpretation of prediction

According to the sequencing principle of ChIP-seq, as we know, this technology is capable of detecting TF-specific binding regions under a certain condition, and these regions are mostly located in open (active) chromatin. However, a large number of such TF binding regions are also located in repressive or poised chromatin. Perhaps for this reason, there are still many negative sequences (not peaks) incorrectly predicted by NLDNN as positive sequences (peaks). By calculating the proportion of cell-type-specific TF binding peaks to total peaks that are integrated from ReMap2022, we find that cell-type-specific TF binding peaks account for only a very small proportion of the overall population (Supplementary Fig.6), which suggests that computational methods have difficulty distinguishing active binding regions from repressive or poised regions by relying only on sequence features. To further explain this phenomenon, we selected all false positives (FPs) from chromosome 1 and performed intersection with true peaks, observing that most of FPs are supported by true peaks (Fig.4a). Moreover, as shown in Supplementary Fig.7, a large number of true peaks were incorrectly predicted as false negatives (FNs). To alleviate these issues, we conducted an additional experiment by integrating DNA sequences and chromatin accessibility signals as inputs. By including or excluding chromatin accessibility signals (Fig.4b-d), we find that the performance of NLDNN+CA using accessibility signals is much better than that of NLDNN without accessibility signals, with average performance gains of 18% and 8% in terms of the PRAUC and Pearson correlation, demonstrating that chromatin accessibility signals can enhance the predictive performance by correcting a certain number of FPs and FNs.

**Fig. 4.**
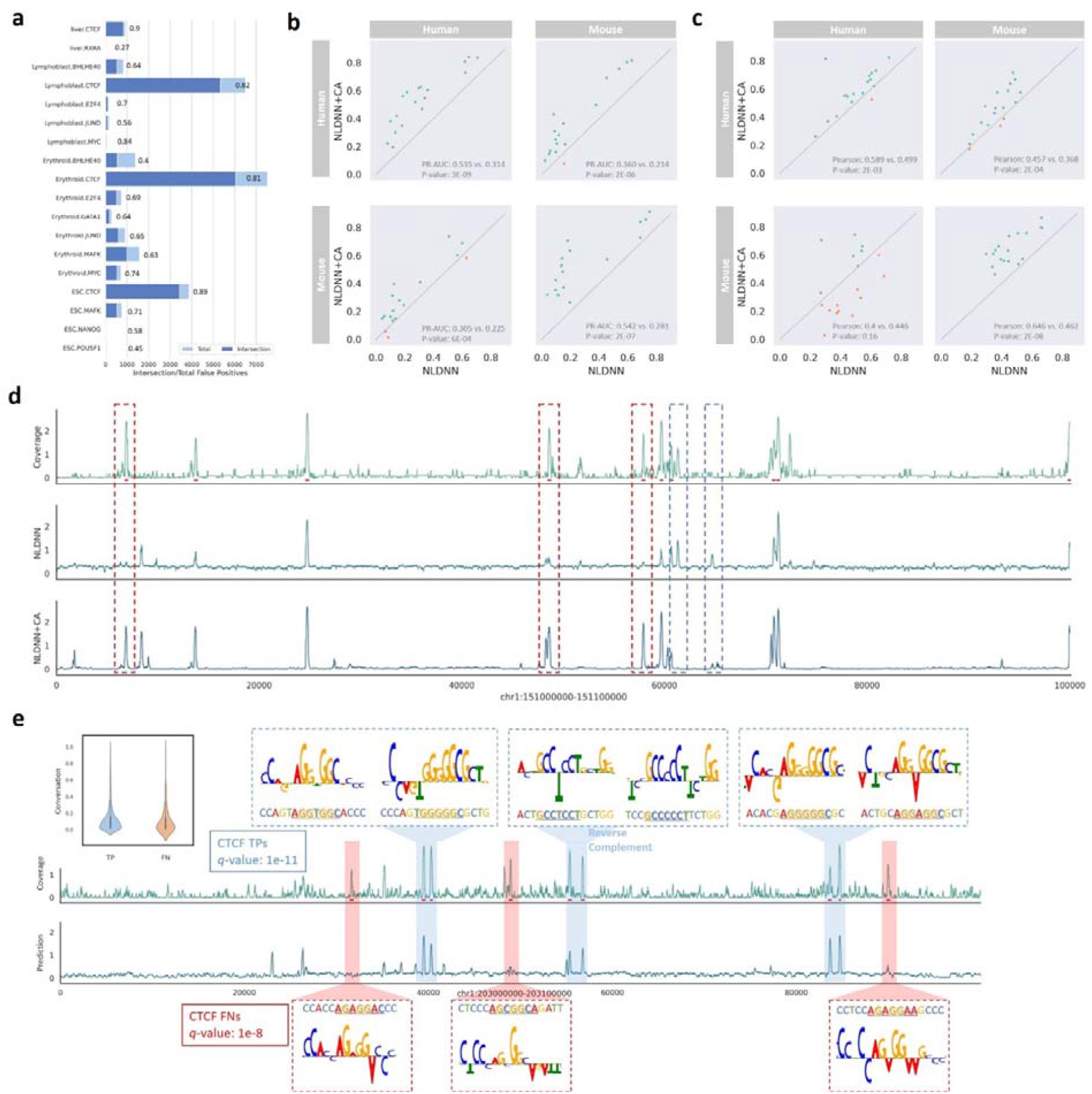
Interpretation of prediction. (a) The proportion of false positives is supported by true peaks. (b-c) Within- and cross-species performance comparison between NLDNN and NLDNN+CA for predicting TF binding, where ‘NLDNN+CA’ means NLDNN integrating DNA sequences and chromatin accessibility (*p*-value is calculated using T-test, one-tailed, paired). (d) Visualization of true coverage values of a human genomic region (chr1:151000000-151100000, 100kb) and corresponding predicted values obtained through NLDNN and NLDNN+CA, where red (blue) dashed boxes represent those false negatives (positives) that are mispredicted by NLDNN but corrected by NLDNN+CA, and red underlines denote true peaks on the genomic region. (e) Visualization of some true positive (false negative) sequences from CTCF binding regions, represented by blue (red) color, respectively. The conservation values were calculated by averaging values extracted from PhastCons100way (UCSC).

Furthermore, we selected true positives (TPs) and false negatives (FNs) from chromosome 1 and performed motif discovery, respectively. An example for CTCF, the matching significance of the CTCF motif learned from TPs is higher than that learned from FNs (*q*-value: 1e-11 vs. 1e-8), implying that TPs contain more conservative motif information than FNs. By application of DeepLift and ISSM, we find that TPs have a conserved short sequence “AGG^*^GGC” while NFs exhibit greater flexibility in conserved sequences, perhaps resulting in low predicted signals for FNs (Fig.4e). The underlying reason is that NLDNN is apt to capture conservative information and thereby assigns high signals to TPs. Another example for GATA1, the significant motif learned from TPs is GATA1 (*q*-value: 1e-9) while the significant one learned from FNs is GATA4 (*q*-value: 1e-3). By applying DeepLift and ISSM, we find that TPs have GATA1-like motifs with high ISSM values while FNs have GATA4-like motifs with high ISSM values (Supplementary Fig.8).

### Reduced cross-species predictive performance

Cross-species TF binding prediction is complicated by the rapid evolutionary turnover of individual TF binding sites across the genomes of different species. This complexity could be reflected quantitatively by the fact that the performance of within-species prediction is evidently better than that of cross-species prediction over all datasets (Fig.5a). By visualizing the coverage values of human DNA regions predicted by human and mouse models (Supplementary Fig.9), intuitively, we observe that a few peaks correctly predicted by human models (high values) are incorrectly predicted by mouse models (low values). Although human and mouse motifs (represented by PWMs) are highly similar, e.g., 99% similarity (Pearson correlation) for the CTCF motif, cross-species TF binding prediction is still influenced by several potential factors, such as repeat elements and flanking sequences. Repeat elements, such as *Alu* elements, a type of SINE, cover a large portion (10%) of the human genome[27], have been systematically investigated by Cochran *et al*.[23], which concludes that reduced cross-species predictive performance can be attributed to one type of species-specific repeats. Inspired by this work, we constructed non-SINE models for both human and mouse species by filtering out any training windows that intersect with *Alu* elements and then evaluated them using all test windows that include *Alu* elements (Supplementary Note 1). From the comparison (Fig.5b), we reveal that in most cases, the predictive performance of non-SINE models is worse than that of models trained using complete windows, indicating that repeat elements indeed play a particular role in predicting TF binding within or across species.

**Fig. 5.**
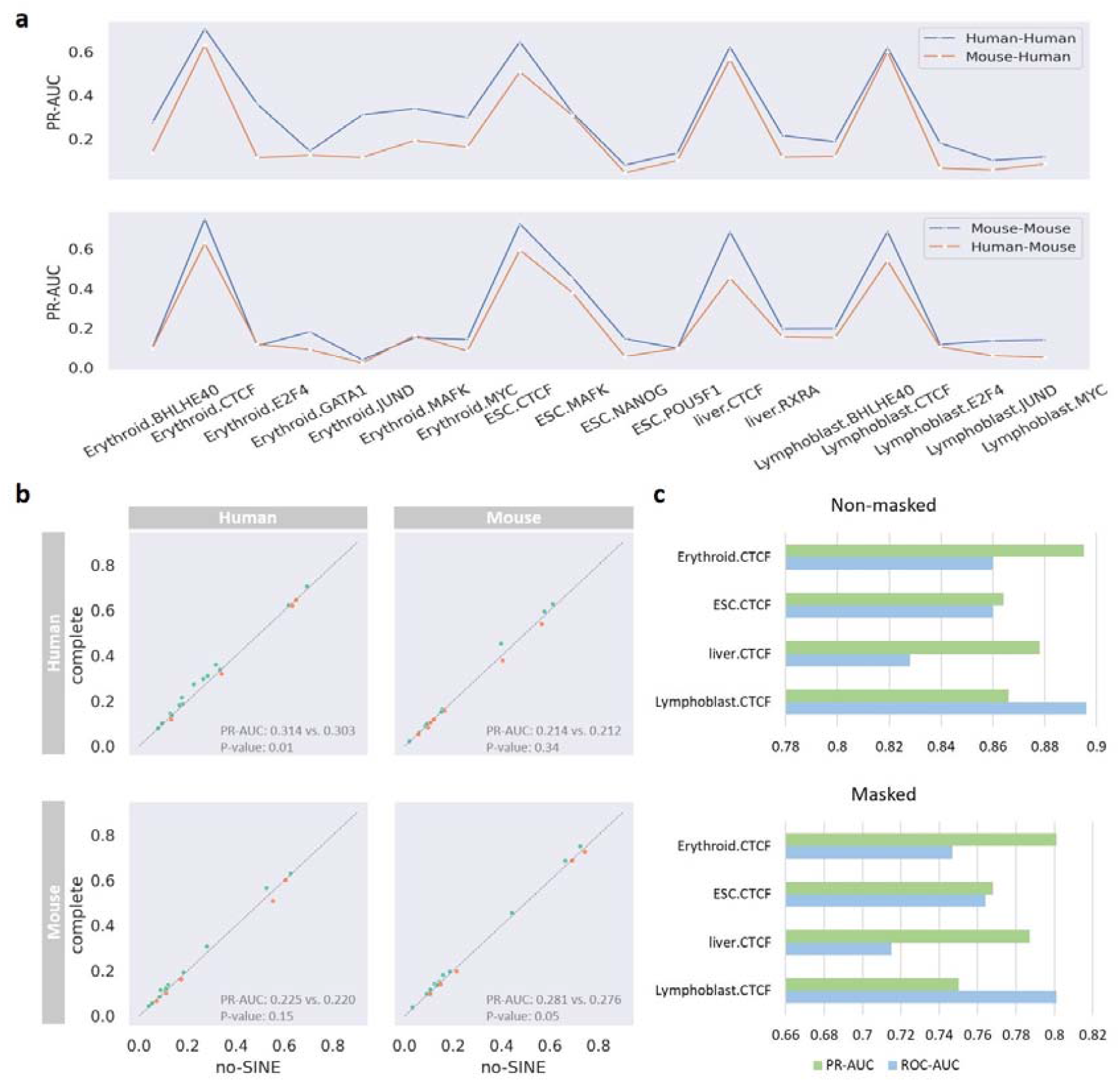
Reduced cross-species predictive performance. (a) Predictive performance comparison between within- and cross-species, where blue (yellow) line means within (cross) species prediction. (b) Within- and cross-species performance comparison between no-SINE and complete models for predicting TF binding (*p*-value is calculated using T-test, one-tailed, paired). (c) Performance of models for discriminating cross-species CTCF binding peaks, where ‘Masked’ means masking CTCF motif instances with high significance from all binding sequences.

As mentioned above, human and mouse motifs are highly similar, meaning that their TF binding sites are similar, but the sequence context features (flanking sequences) of binding sites are predictive of TF binding[28]. To investigate the effect of sequence context features across species, we designed two experiments in which only the sequence context features were used to discriminate cross-species CTCF binding peaks (Supplementary Note 2). According to the results (Fig.5c), we observe whether CTCF binding sites are eliminated or not, CTCF binding peaks for human and mouse are easily distinguishable by relying on the sequence context features of binding sites. This observation demonstrates that the context features of binding sites between human and mouse are species-specific although their binding sites are similar, complicating cross-species prediction of TF binding.

### Adversarial training can improve the performance of cross-species TF binding prediction

As mentioned above, cross-species TF binding prediction is confused by several factors, especially repeat elements, resulting in reduced cross-species predictive performance. To bridge this gap and reduce the difference in cross-species performance, Cochran et al. proposed an unsupervised domain adaptation method that utilizes a gradient reversal layer (GRL) to encourage the “feature generator” portion of a neural network to learn species-generic features[29]. Although this method performs well on cross-species prediction of four TFs in liver, it fails to generalize well to an expanded set of TFs from different cell lines. Recent studies[30, 31] have revealed that adversarial learning from Generative Adversarial Networks (GAN) has advantages over GRL in the field of domain adaptation. Given this point, we design a dual-path framework to mimic the framework of GAN in which a source generator is to generate feature mappings from a source species (‘real’ data), a target generator is to generate feature mappings from a target species (‘fake’ data), and a discriminator is to discriminate the ‘real’ and ‘fake’ data. Correspondingly, NLDNN is required to be decomposed into a generator and a predictor in which the generator is fine-tuned by adversarial learning while the predictor is directly appended onto the fine-tuned generator to predict cross-species coverage values (see **Methods** for details). Initially, in the process of constructing data, we randomly sampled sequences from whole genomes of human and mouse species, respectively, and then used these data to fine-tune the target generator by adversarial training in the dual-path framework. Compared with NLDNN, the performance of cross-species prediction is not improved across all TF binding datasets (Supplementary Fig.10a). We speculate that the underlying reason may be the lack of TF binding information in the sampled data. Therefore, binding sequences from the target species were gradually added into sampled data in an increasing proportion, e.g., {0, 0.001, 0.1, 0.5, 1}. Similarly, we evaluated NLDNN-AT using three criteria: (i) predictive performance, (ii) variant effect prediction, and (iii) localization ability.

To demonstrate the overall performance of NLDNN-AT for predicting cross-species TF binding, we compared it with three competing methods across 18 TF binding datasets (Fig.6a-b). Intuitively, NLDNN-AT outperforms NLDNN and NLDNN-TL in terms of classification (PR-AUC) and fitting (Pearson correlation) performance. Moreover, GRL performs worse than NLDNN-AT although it surpasses its basic framework (DanQV). These observations demonstrate the effectiveness of the dual-path framework of adversarial training for cross-species prediction. Beyond predictive performance, we also made a comparison of NLDNN and NLDNN-AT for variant effect prediction and localization ability. For variant effect prediction (Fig.6c-d), in most cases, NLDNN-AT performs better than NLDNN for classifying filtered or non-filtered TF-specific SNPs. For localization ability (Fig.6e-f), according to the number of located regions intersecting with true peaks, NLDNN-AT is generally better than NLDNN for locating true peaks, especially for mouse-to-human. In addition, we used the dual-path framework to fine-tune NLDNN+CA that incorporates accessibility signals by adversarial training, finding that the cross-species predictive performance of NLDNN-AT+CA is evidently better than that of NLDNN+CA in terms of PR-AUC and Pearson correlation (Fig.6a-b). The above results fully demonstrate that adversarial training can improve the performance of cross-species TF binding prediction. To better understand how adversarial training improves predictive performance, we visualized the coverage values of human DNA regions separately predicted by mouse and mouse-adaptation models (Fig.6g and Supplementary Fig.9). As a result, we can directly see that mouse-adaptation models pay attention to all potential peaks found by mouse models and amplify the predicted coverage values of them so that some peaks incorrectly predicted by mouse models are corrected by mouse-adaptation models.

**Fig. 6.**
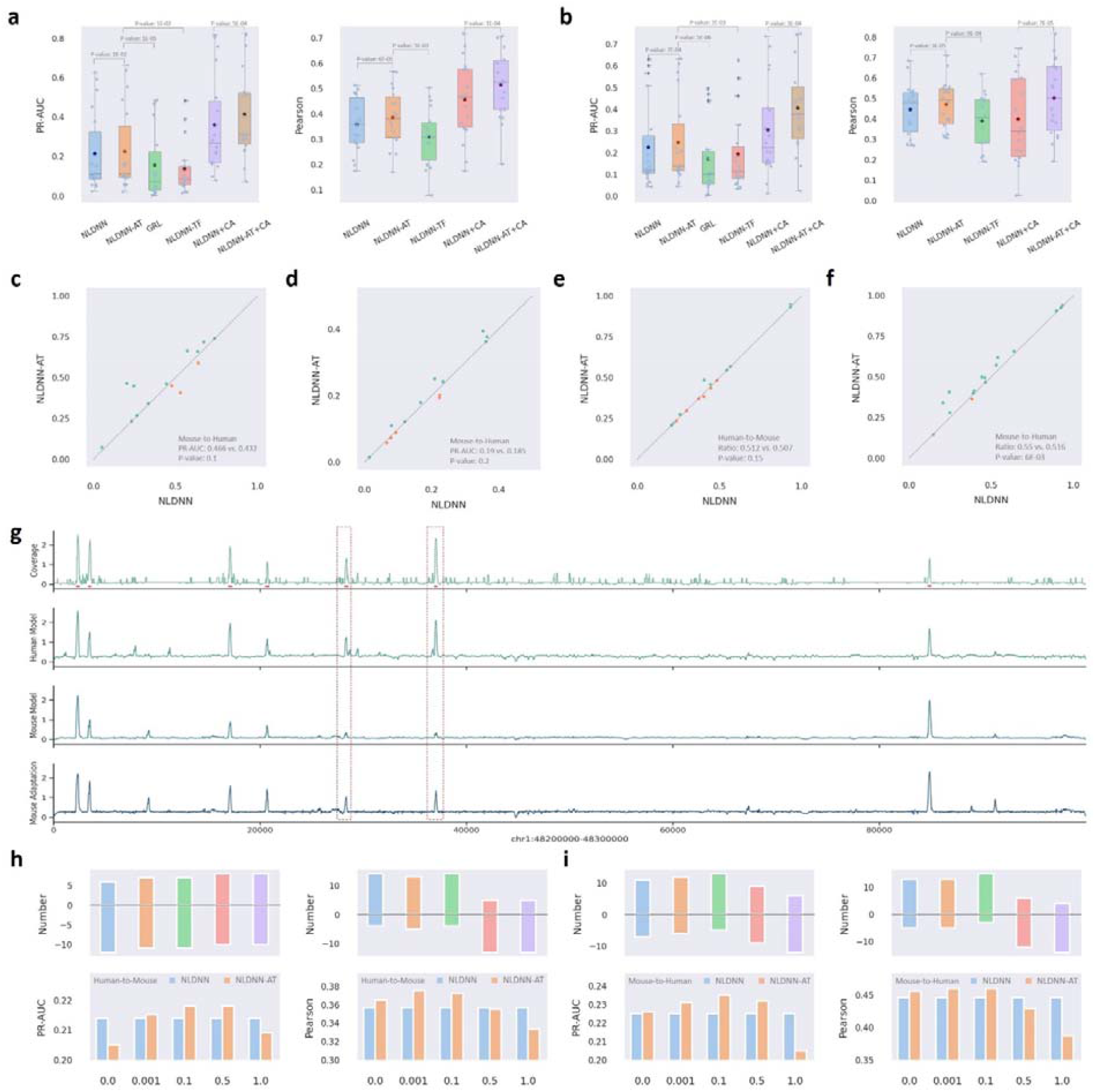
Overall cross-species predictive performance of NLDNN-AT. (a) Cross-species performance (Human-to-Mouse) comparison between NLDNN-TF and competing methods for predicting TF binding. (b) Cross-species performance (Mouse-to-Human) comparison between NLDNN-TF and competing methods for predicting TF binding. (c) Performance comparison between NLDNN and NLDNN-AT for classifying TF-specific SNPs filtered by peaks. (d) Performance comparison between NLDNN and NLDNN-AT for classifying TF-specific SNPs not filtered by peaks. (e) Cross-species performance (Human-to-Mouse) comparison between NLDNN and NLDNN-AT for locating TF binding regions. (f) Cross-species performance (Mouse-to-Human) comparison between NLDNN and NLDNN-AT for locating TF binding regions. (g) Visualization of true coverage values of a human genomic region (chr1:48200000-48300000, 100kb) and corresponding predicted values obtained through human, mouse, and mouse-adaptation models, where red dashed boxes represent those peaks that are mispredicted by mouse models but corrected by mouse-adaptation models, and red underlines denote true peaks on the genomic region. (h) Cross-species predictive performance (Human-to-Mouse) of NLDNN-AT under different proportions of binding sequences from the mouse species. (i) Cross-species predictive performance (Mouse-to-Human) of NLDNN-AT under different proportions of binding sequences from the human species. *p*-value is calculated using T-test, one-tailed, paired.

To investigate the contribution of TF binding information, the results of NLDNN-AT were decomposed into five parts with different proportions of binding sequences. In addition to the PR-AUC and Pearson correlation, we evaluated the number of TF binding datasets that exhibited improved performance. According to the three metrics, for human-to-mouse prediction (Fig.6h), models with low proportions (0, 0.001, 0.1) are generally better than those with high proportions (0.5, 1); for mouse-to-human prediction (Fig.6i), there is a more significant improvement in performance, and models with low proportions (0, 0.001, 0.1) are better than those with high proportions (0.5, 1). More evidently, models that use all binding sequences from the target species perform worst for cross-species prediction. These observations imply that TF binding information and background information of species complement each other and jointly contribute to cross-species prediction. In the same way, the results of NLDNN-AT+CA were decomposed into five parts with different proportions of binding sequences. Instead, models with high proportions are generally better than those with low proportions under the influence of chromatin accessibility, indicating that chromatin accessibility representing all opening regions needs more TF binding information to assist cross-species prediction (Supplementary Fig.10b).

## Discussion

Although NLDNN works well on the task of TF binding prediction, it fails to surpass sequence-level models (DeepSea and DeepSea+) on the task of chromatin accessibility prediction. The former task focuses on TF-specific binding patterns while the latter focuses on TF-generic binding patterns, implying that nucleotide-level models are good at mining TF-specific motif features while sequence-level models are good at integrating the context information of whole sequences. In the experiments, the low classification performance of NLDNN (PR-AUC < 0.3) may be caused by the ways of selecting negative sequences; for instance, in this study, we used the whole genome excluding binding peaks as negative sequences. Obviously, this selection will make TF binding prediction more difficult, as negative sequences will contain a large number of inactive or poised regions bound by the same TF or regions bound by the same TF family. Instead, we adopted a simple way to select negative sequences, such as the upstream sequences of binding peaks, by which the vast majority of confusing regions are eliminated, thereby getting a very high PR-AUC value (Supplementary Fig.11a). This observation makes us understand that the selection of negative samples is closely related to the final prediction and rethink how to elaborate negative samples. In addition, we cut binding peaks into sequences of 600bp length in this study; however, dilated convolution (BPNet) has a wider receptive field to take advantage of longer sequences. To investigate it, we expanded the length of sequences from 600bp to 2000bp and re-trained NLDNN and BPNet on these 2000bp sequences. As a result, we find that longer sequences impose a positive impact on classification performance but a negative impact on fitting performance (Supplementary Fig.11b), demonstrating that the extended contextual information of binding sites is beneficial for discriminating binding and non-binding sequences but may bring noise to regression of coverage values.

As shown in Supplementary Fig.9, although NLDNN-AT is able to correct some mispredictions by amplifying the coverage values of incorrectly predicted peaks, it enhances the predictions of all potential peaks and cannot recognize whether they belong to TPs or not, indicating that adversarial training is unable to reduce the number of FPs and completely dependent on the results produced by its basic framework, such as NLDNN. A possible solution is to utilize more contextual information of sequences. However, although NLDNN employs GRU to capture the long-term dependencies in sequences, GRU may encounter gradient vanishing or exploding problems when dealing with long sequences. Emerging architectures based on transformer[32] could utilize the self-attention mechanism to better capture longer-range dependencies in sequences, without being affected by the gradient vanishing or exploding problems of GRU. Moreover, the framework of adversarial training can easily be transferred to other architectures. Thus, a combination of transformer-based architectures and adversarial training could provide a promising solution for reducing FPs in cross-species prediction.

## Conclusions

In recent years, the number and variety of DL models for predicting regulatory genomic tasks have been continuously increasing, which are roughly divided into two categories: sequence-level models and nucleotide-level models. In contrast, nucleotide-level models can provide a nucleotide-level perspective for studying TF binding and its related tasks. Accordingly, we develop a novel Nucleotide-Level Deep Neural Network (NLDNN) to predict TF binding within and across species. However, the question of how to directly compare sequence-level and nucleotide-level models remains unsolved. To solve this question, we propose to use the maximums of coverage values as a bridge to make a direct comparison of the two types of models, by which we can calculate two quantitative indicators, such as PR-UAC and Pearson correlation. Experimental results show that NLDNN performs best for predicting TF binding within and across species in terms of predictive performance, variant effect prediction, and localization ability. Besides, we make a comprehensive analysis of the contributions of nucleotides and explore the effect of variants on TF binding prediction by model interpretability and ISSM. To further improve the performance of predicting cross-species TF binding, inspired by GAN, we design a dual-path framework to fine-tune NLDNN by adversarial training (NLDNN-AT). Through experiments and analysis, we conclude that adversarial training can improve the performance of cross-species TF binding prediction by amplifying the predicted coverage values so that some peaks incorrectly predicted by mouse models are corrected by mouse-adaptation models.

## Methods

### Data processing

ChIP-seq data for human (hg38) and mouse (mm10) were acquired from the Encyclopedia of DNA Elements (ENCODE) and the National Center for Biotechnology Information (NCBI), including 18 TFs from 4 cell lines/types that were previously used by Cochran, et al., each of which contains an irreproducible discovery rate (IDR) peak file and a nucleotide-resolution coverage track file. The nucleotide-resolution coverage values are expressed in two ways: fold-over control at each position, or p-value to reject the null hypothesis that the value at that location is present in the control. In the subsequent analysis, we found that p-value tracks are better suitable for our proposed method than fold change over control tracks, so p-value coverage tracks were selected as nucleotide-resolution supervision signals (Supplementary Fig.11c). Experimental accessions for all data used in this study are listed in Supplementary Table 1.

The human (hg38) and mouse (mm10) genomes were split into 600bp windows, offset by 100bp. Any windows overlapping ENCODE blacklist regions were removed. For each ChIP-seq data, positive sequences were constructed by selecting windows whose ratio overlapping with the IDR peak exceeds 0.2, while negative sequences were constructed by using all windows that do not overlap with the IDR peak and match the GC distribution of positive sequences. Then, nucleotides of each window were transformed into a matrix of 4^*^600 where ‘4’ represents four types of nucleotide {A, C, G, T}. Correspondingly, nucleotide-resolution coverage values for positive sequences were extracted from the coverage track file and scaled by the log_10_ function, while nucleotide-resolution ‘zero’ values were generated for negative ones.

Chromosomes 1, 18, and 8 of both species were held out from all training data (excluding Chromosome Y) where Chromosome 8 was used as the validation set for hyper-parameter tuning while Chromosome 1 and 18 were used as the test set.

## Models

### NLDNN

In this paper, we propose a novel Nucleotide-Level Deep Neural Network (NLDNN) to predict TF binding within and across species. NLDNN takes DNA sequences as input and directly predicts experimental coverage values as output, which regards TF binding prediction as a nucleotide-level regression task. Inspired by U-Net[24], a typical model for image semantic segmentation, NLDNN is composed of an encoder architecture for gradually reducing the resolution of inputs and learning sequence-specific features, a decoder architecture for gradually enlarging the resolution of features and predicting the coverage values of nucleotides, and a connection architecture for integrating the information flow of the encoder and decoder. The architectures of NLDNN are depicted in Fig.1a-b.

### Encoder architecture

It contains three convolutional blocks, a bi-directional GRU (Gated Recurrent Unit[33]) layer, and a SPPM (Simple Pyramid Pooling Module[34]) layer, in which each convolutional block is made up of a convolutional layer, an ELU (Exponential Linear Unit[35]) layer, a max-pooling layer as well as a dropout layer. Specifically, the convolutional block is used to gradually reduce the resolution of inputs and encode sequence-specific features; the bi-directional GRU layer is used to learn the long-term dependencies between sequence-specific features; the SPPM layer is used to capture the multi-scale context of genomic sequences.

### Decoder architecture

It contains three up-sample blocks and an output layer, in which each up-sample block is composed of an up-sample layer and a blending layer that consists of a batch normalization (BN) layer, a ReLU layer, and a convolutional layer. Specifically, the up-sample layer is used to enlarge the resolution of down-sampled features; the blending layer is used to re-adjust the values of up-sampled features; the output layer is used to predict the coverage values of nucleotides.

### Connection architecture

It contains three RRBs (Refinement Residual Block[36]), and each RRB is used to integrate the same-level information flow of the encoder and decoder.

The detailed parameter settings of NLDNN are illustrated in Supplementary Fig.12.

### Sequence-level models

These models mainly focus on prediction tasks at the sequence level, including LSGKM[37], DeepSea[6], DanQ[11], and a variant of DanQ (DanQV[23]). Specifically, LSGKM is a new version of gkm-SVM for large-scale datasets, which offers much better scalability and provides further advanced gapped *k*-mer based kernel functions for detected functional regulatory elements in DNA sequences. DeepSea is composed of three convolutional layers (each followed by a ReLU layer, a max-pooling layer, and a dropout layer) and two fully-connected layers. DanQ is composed of a convolutional layer followed by a ReLU layer, a max-pooling layer, a dropout layer, a bi-directional LSTM (Long Short-term Memory) layer followed by a dropout layer, and two fully-connected layers. DanQV is composed of a convolutional layer followed by a ReLU layer, a max-pooling layer, a LSTM (Long Short-term Memory) layer, and two fully-connected layers. Note that, except for LSGKM, a sigmoid layer is added to the end of these models for binary classification, otherwise for regression. For classification models, “1” was used to label positive sequences, while “0” was used to label negative sequences. For regression models, the maximums of experimental coverage values were used to label positive sequences, while “0” was used to label negative sequences.

### Nucleotide-level models

These models mainly focus on prediction tasks at the nucleotide level, including two nucleotide-level regression models (BPNet[14] and FCNsignal[15]) and two nucleotide-level classification models (Leopard[13] and FCNA[12]). BPNet consists of a convolutional layer, followed by nine dilated convolutional blocks with progressively increasing dilation rates (scaled by powers of 2) in which each has a residual connection to the previous layer. It should be noted that a key difference from the original BPNet is that the negative strand, bias track, and read counts output head were not used throughout this study. FCNsignal consists of three convolutional blocks, three up-sampling blocks, and three connection lines (summation). A key difference from NLDNN is that this method does not incorporate SPPM and RRB. Leopard consists of an encoder containing five convolution-convolution-pooling (CCP) blocks, a decoder containing five upscaling-convolution-convolution (UCC) blocks, and five connection lines (concatenation). It should be noted that a key difference from the original Leopard is that DNase-seq signals were not used throughout this study. FCNA consists of three convolutional blocks, three up-sampling blocks, and four connection lines (summation). A key difference from NLDNN is that this method does not incorporate bi-directional GRU, SPPM, and RRB. Since Leopard and FCNA are nucleotide-level classification models, the last sigmoid layer was removed from their architectures for predicting coverage values.

### Model training

Due to the extreme imbalance between the number of positive and negative sequences, we adopted a ‘random sampling without replacement’ strategy to deal with negative sequences. Specifically, in each training epoch, we used all positive sequences and randomly sampled three times the number of negative sequences without replacement, aiming at iterating over all negative sequences and minimizing the impact of imbalance on model performance. MSE (Mean Squared Error) loss at nucleotide resolution was used to train nucleotide-level models (Equation 1). Except for LSGKM, BCE (Binary Cross Entropy) and MSE loss were separately used to train sequence-level classification and regression models.

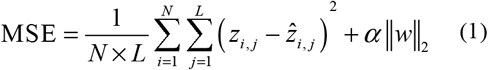

where *N* is the number of samples; *L* is the length of each sequence; *zi, j* and 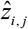 are the predicted and true coverage values, respectively; *α* is a regularization parameter to leverage the trade-off between the goal of fitting and the goal of the generalizability of the trained model; || ||_2_ indicates the L2 norm.

Given that DL-based methods are sensitive to parameter initialization, which may yield significantly different results, we employed a ‘warm-up’ strategy to reduce the impact of parameter initialization on DL-based methods. Specifically, we first warmed up DL-based methods by running some randomly initialized models a few times and selected the best-initialized model in terms of validation accuracy; then, the selected model was used as an initialized template for the training phase. All models were trained with a batch size of 500 using ADAM[38] with default parameters. The number of training epochs was set to 60, and the initial learning rate was set to 0.001 and decayed by a factor of 0.9 every 10 epochs.

### Maximums of coverage values

In this study, we defined the strength of TF binding as the number of reads falling into IDR peaks. To achieve this, we first collected filtered alignments (bam files) from ENCODE and then used the *bedtools* tool to compute the number of reads falling into each peak. The strength values for all peaks were further scaled by the log10 function.

Up to now, a variety of DL-based methods have been proposed for predicting TF binding, roughly categorized as sequence-level models and nucleotide-level models. However, it remains challenging to make a direct comparison of their model performance since the two types of methods have different types of outputs. For example, sequence-level models often output a scalar representing a label probability or regression value while nucleotide-level models often output a vector representing label probabilities or regression values for all nucleotides. To address this problem, we propose to use the maximums of coverage values to represent the strength of TF binding, by which positive (binding) sequences can be distinguished from negative (non-binding) ones since the binding strength of positive sequences is generally higher than that of negative ones. Therefore, a direct comparison of their model performance was realized through the maximums of coverage values.

### Evaluation

To evaluate the overall performance of all models for predicting TF binding within and across species, PR-AUC (Area Under the Precision-Recall Curve) was adopted to verify the classification performance of all models. Specifically, (i) for sequence-level classification models, the predicted probabilities and true labels (0/1) were used to calculate the PR-AUC; (ii) for sequence-level regression models, the predicted and true maximums were used to calculate the PR-AUC; (iii) for nucleotide-level models, the maximums of predicted and true coverage values were used to calculate the PR-AUC.

Pearson correlation was adopted to verify the fitting (regression) performance of all models on positive sequences. Specifically, (i) for sequence-level classification models, the predicted logits (after removing sigmoid) and true maximums were used to calculate the Pearson correlation; (ii) for sequence-level regression models, the predicted and true maximums were used to calculate the Pearson correlation; (iii) for nucleotide-level models, the maximums of predicted and true coverage values were used to calculate the Pearson correlation. Besides, the Pearson correlation between the predicted and true coverage values of all nucleotides was also calculated for nucleotide-level models. The full evaluation process is shown in Fig.2a.

### Variant effect prediction

#### TF-specific SNPs

To construct TF-specific SNPs, (i) we collected eQTL SNPs from the GTEx project[39], targeting at Liver, EBV-transformed lymphocytes, and Whole Blood which are related to cell lines used in this study; (ii) we collected TF-specific ChIP-seq peaks from ReMap2022 and corresponding TF PWMs from HOCOMOCO[40], and then utilized FIMO[41] to find potential motif instances with high significance (p-value < 1e-4) on these peaks; (iii) we picked out SNPs that are located in the region of any motif instances as TF-specific SNPs. To create a binary classification task, we regarded TF-specific SNPs as the positive set and selected all neighboring SNPs around each TF-specific SNP (distance < 5kb) from dbSNP[42] as the negative set, based on an assumption that mutation sites in adjacent regions of genomes have a certain genetic correlation, called Linkage Disequilibrium (LD)[43]. To get convincible results, only Chromosomes 1 and 18 were used to construct TF-specific SNPs since both are not observed by trained models. Besides, we also constructed non-filtered TF-specific SNPs using the same process, except that we utilized FIMO to find potential motif instances with high significance (p-value < 1e-4) on a whole genome. For each SNP, 600bp DNA sequences centered on reference and alternative alleles were separately extracted and then transformed into one-hot matrices.

#### MPRA data

This data consists of massively parallel reporter assays (MPRA) that measure the effect size of all single-nucleotide variants through saturation mutagenesis of several regulatory elements across different cell types[43], 15 of which were used for variant effect prediction in this study. For each variant, 600bp DNA sequences centered on reference and alternative nucleotides were separately extracted and then transformed into one-hot matrices.

#### Causal SNPs

Three disease-associated SNP groups with strong LD effects were collected. Candidate causal SNPs of myeloma consisting of 10 risk variants at 7p15.3 that alter IRF4 binding[44], pan-autoimmune consisting of seven genetic susceptibility variants at 6q23 that influence the binding of eight TFs[45], and chronic lymphocytic leukemia (CLL) consisting of 27 risk variants at 15q15.1 that disrupt the binding of RELA[46] were obtained from previous literature. For each SNP, 600bp DNA sequences centered on reference and alternative alleles were separately extracted and then transformed into one-hot matrices.

### Model preparation

For TF-specific SNPs, (i) TF-specific models were trained using the above-described TF binding data; (ii) the trained models took one-hot matrices of SNPs as input and predicted the label probabilities (for sequence-level models) or coverage values (for nucleotide-level models); (iii) since we only focused on the classification performance, we defined the effect values as the absolute values of the difference of outputs between alternative and reference sequences, e.g., |*p*^*alt*^ − *p*^*ref*^| for sequence-level models, and 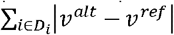 for nucleotide-level models where *i* denotes the position of SNPs and *D*_*i*_ the neighboring region of position *i*. For MPRA data, (i) we downloaded related DNase- or ATAC-seq data from ENCODE and processed it using the same way described above, except that the chromosomes containing these SNPs were used, as the test set while the remaining chromosomes were used as the training set; (ii) the trained models took one-hot matrices of SNPs as input and predicted the label probabilities or coverage values; (iii) we defined the effect value as palt - pref for sequence-level models and 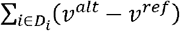) for nucleotide-level models.For causal SNPs, we downloaded GM12878 ATAC-seq data from ENCODE that are related to these SNPs, and used the same process as the one for MPRA data to prepare models.

### Evaluation

For TF-specific SNPs, ROCAUC and PRAUC were used to evaluate the performance of models for distinguishing positive SNPs from negative SNPs. For MPRA data, Pearson correlation and Spearman correlation were used to evaluate the performance of models for fitting MPRA data. For causal SNPs, their effect values were directly used to prioritize these SNPs, of which the best effect value was regarded as the causal one associated with diseases. Then, the number of correctly matched ones was used to evaluate the performance of models for prioritizing real causal SNPs. Besides, the ratio of the effect value of the real SNP to the highest value of the remaining SNPs with strong LD was calculated to assess the ability to discriminate causal SNPs.

### Locating potential TF binding regions

#### Motivation

Given that the maximums of coverage values are most likely located in positive (binding) sequences, so the predicted maximums were used to locate all potential TF binding regions in this study. Specifically, Chromosome 1 from the test set was first segmented into non-overlapping windows of 600bp, and the trained models took windows as input and predicted their coverage values. Then, the positions of the maximums of coverage values were mapped to the genome. Finally, we can manually set a high threshold value to filter out negative windows or select the top 1% of windows as potential TF binding regions. To make a fair comparison of nucleotide-level models, we chose the second way to select potential TF binding regions which can guarantee that the total number of selected binding regions is consistent.

#### Evaluation

To evaluate the performance of nucleotide-level models for locating TF binding regions, we designed two ways: a direct way and an indirect way. With regard to the direct way, (i) TF ChIP-seq peaks were downloaded from ReMap2022[47], which were integrated from different cell lines; (ii) the positions of all potential TF binding regions (top 1%) were used to intersect with TF ChIP-seq peaks, counting 1 if intersected, otherwise 0; (iii) the ratio of intersected number to total number was used as a quantification indicator of evaluation. With regard to the indirect way, (i) corresponding TF PWMs were collected from HOCOMOCO[40]; (ii) FIMO[41] coupled with PWMs was run on all potential TF binding regions to find motif instances with high significance (*p*-value < 1e-4); (iii) the ratio of the number of motif instances to total number was used as another quantification indicator of evaluation.

### Model interpretability

#### Model itself

We can utilize the model itself to implement the task of motif discovery. For a given NLDNN model, for example, (i) a trained model was used to predict the coverage values of each test sequence; (ii) the position of the maximum of the predicted values was located in the sequence, and then a window of length 60bp surrounding this position was extracted; (iii) the first convolutional layer was used to score each sub-region of the window by a stride size of 1 and then a sub-region with the highest score was selected; (iv) all selected sub-regions from the test set were aligned to compute corresponding PWMs (each kernel corresponds to a PWM); (v) TOMTOM[48] was employed to match the learned PWMs with experimentally validated motifs from standard databases, e.g, HOCOMOCO. Subsequently, those matched motifs with high *q*-value that do not belong to the same TF family as the target motif were regarded as potential co-binding motifs.

#### Contribution scores

Contribution scores of nucleotides were computed by DeepLIFT[49], which is a feature attribution method for computing the contribution of each nucleotide (feature) in an input sequence to a specific scalar output prediction from a neural network model. DeepLIFT decomposed the difference between the prediction from an input sequence versus that of a neutral reference sequence. As suggested by relevant literature, all zeroes were used for reference sequences. However, the outputs of nucleotide-level models are not a scalar but a vector of size *L* where *L* is the sequence length. Therefore, for adapting DeepLIFT, we need to adjust nucleotide-level models by appending a max function to get a scalar (maximum) since we are mainly concentrated on the maximum of coverage values. For a given model, a contribution matrix of size *A* ^*^ *L* was obtained through DeepLIFT, where *L* is the sequence length and *A* is 4 (one for each nucleotide). Then, the matrix was multiplied by the input sequence (one-hot matrix) to obtain the sensitivity of the observed nucleotide at each position.

#### TF–Modisco

TF–Modisco[50] was run on the contribution scores of the test set profiled by DeepLIFT for each TF, respectively. Significant seqlets were selected by computing contribution scores over a width of 21 bp and using the false-discovery rate threshold of 0.01. CWMs (Contribution Weight Matrice) were then computed from the aligned seqlets by averaging the contribution scores. At last, TOMTOM was employed to match the learned CWMs with experimentally validated motifs from standard databases.

#### Evaluation

To evaluate the matching degree between learned motifs and validated motifs, a significance indicator *q*-value was used and then transformed by the – log_10_ function.

### In silico saturation mutagenesis

In silico saturation mutagenesis (ISSM) of an input sequence was performed by calculating the effect values of variants at each position in which each variant is generated by replacing the original nucleotide with the remaining nucleotides. For example, the *i*^th^ position’s nucleotide in a 600bp sequence is A, and corresponding variants at that position are A→ C, A→ G, and A→ T, respectively. Then, the effect values of these variants are calculated according to the equation 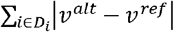 where *D*_*i*_ denotes the neighboring region of the position *i*. As a result, a scoring matrix of size 4 ^*^ 600 is constructed where the element 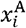 in the matrix represents the effect value of the reference nucleotide mutated to the nucleotide A at position *i*, and the larger the value of a variant, the greater its importance for TF binding.

### Categorization of predicted binding sites

In order to categorize the predicted sites as false positives (FPs), true positives (TPs) and false negatives (FNs), we adopted the following steps: (i) the predicted and true coverage values were converted to a range of -1 to 1 by the equation: 2^*^ (*x* − *x*_*min*_)/(*x*_*max*_ − *x*_*min*_) − 1, and then scaled to 0∼1 by the sigmoid function; (ii) we defined binding sites as FPs by the rule: predictions (*P*) are above 0.5 while trues (*T*) are below 0.5 and |P-T| > 0.5, as TPs by the rule: both predictions (*P*) and trues (*T*) are above 0.7, as FNs by the rule: predictions (*P*) are below 0.5 while trues (*T*) are above 0.5 and |*P* − *T*| > 0.5.

### The framework of adversarial training

To further improve the performance of the proposed method NLDNN for predicting cross-species TF binding, we design a dual-path framework and propose an adversarial training strategy inspired by GAN to fine-tune NLDNN by pulling closer the domain space of the human and mouse species.

#### Data construction

DNA sequences of one species were used as the source data labeled as “1” while DNA sequences of another species were used as the target data labeled as “0”, in which DNA sequences of both species are from the training set. Given that the source species is observed and binding sequences are informative of TF binding, binding sequences from the source species were therefore used as the source data. On the contrary, since the target species is unobserved, arbitrary sequences from the target species were used as the target data. In addition, to investigate the contribution of TF binding information, binding sequences from the target species were gradually added into the target data in an increasing proportion, e.g., {0, 0.001, 0.1, 0.5, 1}.

#### Model architectures

As shown in Fig.1a-c, the basic architectures include a generator, a predictor, and a discriminator, in which the generator and predictor are two parts of NLDNN. Specifically, the generator takes DNA sequences as input and generates feature mappings; the predictor takes these feature mappings as input and predicts coverage values; the discriminator takes these feature mappings from the source and target species as input and predicts the label probability of species. The discriminator consists of three convolutional blocks, a BN layer, and two fully-connected layers separately followed by a ReLU layer and a sigmoid layer, in which each convolutional block is composed of a convolutional layer, a ReLU layer, a max-pooling layer as well as a dropout layer.

As shown in Fig.1e, the full framework is a dual-path architecture where the top path contains a source generator for generating feature mappings from the source species while the bottom path contains a target generator for generating feature mappings from the target species, on top of which a discriminator is appended to discriminate the source and target species. Our main goal is to minimize the distance between the distribution of the source and target feature mappings. Therefore, we propose an adversarial training strategy to train the target generator and discriminator iteratively. The training process was performed in multiple stages, more precisely: (i) the source generator was trained by minimizing the MSE loss (Equation 1) over labeled data (including DNA sequences and coverage values) from the source species (Fig.1d), and then the target generator was initialized with the pre-trained source generator; (ii) keeping the target generator unchanged, the discriminator was trained by minimizing the standard binary classification (Equation 2) over the feature mappings generated from the source and target species and corresponding species labels; (iii) keeping the discriminator unchanged, the target generator was trained by minimizing the inverted label GAN loss[51] (Equation 3) over the feature mappings generated from the target species. Note that the source generator is fixed all the time during adversarial training. The full framework was trained using the constructed data with ADAM in which the parameter betas was set to (0.5, 0.9), and the initial learning rates for the generator and discriminator were set to 1e-05 and 1e-04, respectively. After adversarial training, as shown in Fig.1f, the target generator and predictor were concatenated to predict the coverage values of the test set from the target species. For simplicity, the method is abbreviated as NLDNN-AT.

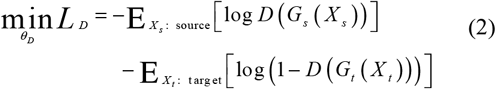

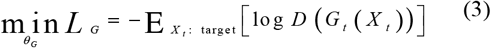

#### Competing methods

A domain-adaptive method for cross-species prediction of TF binding was recently proposed[29], which builds upon the DanQV model described above by adding a new “species discriminator” branch. This method puts a gradient reversal layer (GRL) before the new branch to achieve domain-adaptive training, which automatically discourages learning of training species-specific sequence features by merely outputting the identity of its input during forward propagation but multiplying the gradient of the loss by -1 during backpropagation. This method was trained using the constructed data with default parameters. For simplicity, the method is abbreviated as GRL.

Transfer learning is a commonly-used technique for cross-species prediction tasks[52, 53], we, therefore, applied transfer learning to fine-tune NLDNN for cross-species prediction of TF binding but only using species labels. To achieve this, we added a new classifier on top of the trained source generator, which consists of a global average pooling layer and two fully-connected layers separately followed by a ReLU layer and a sigmoid layer. The full framework was trained using the constructed data with ADAM in which the learning rate was set to 1e-5 for the source generator but 1e-3 for the new classifier. After training, the fine-tuned generator and predictor were concatenated to predict the coverage values of the test set from the target species. For simplicity, the method is abbreviated as NLDNN-TL.

#### Evaluation

To evaluate the overall performance of these methods for predicting cross-species TF binding, PR-AUC was used to evaluate their classification performance, and Pearson correlation was used to evaluate their fitting performance.

## Availability of data and materials

The data accession list can be found in Supplementary Table 1. The source code can be found at http://github.com/turningpoint1988/NLDNN-AT.

## Acknowledgments

This work was supported in part by STI 2030—Major Projects, under Grant 2021ZD0200403, and partly supported by grants from the National Science Foundation of China, Nos. 62333018, 62372255, U22A2039, 62073231, 61932008, and 62372318, and supported by the China Postdoctoral Science Foundation under Grant No.2023M733400, and supported by the Key Project of Science and Technology of Guangxi (Grant no. 2021AB20147), Guangxi Natural Science Foundation (Grant nos. 2021JJA170204 & 2021JJA170199) and Guangxi Science and Technology Base and Talents Special Project (Grant nos. 2021AC19354 & 2021AC19394), and supported by the Natural Science Foundation of Ningbo City under Grant No.2023J199, and supported by Key Research and Development (Digital Twin) Program of Ningbo City under Grant Nos.2023Z219, 2023Z226.

